# The development of 3D bovine intestinal organoid derived models to investigate *Mycobacterium avium* ssp *paratuberculosis* pathogenesis

**DOI:** 10.1101/2022.05.13.491821

**Authors:** Rosemary Blake, Kirsty Jensen, Neil Mabbott, Jayne Hope, Jo Stevens

## Abstract

*Mycobacterium avium* subspecies *paratuberculosis* (MAP) is the etiologic agent of Johne’s Disease, a chronic enteritis of ruminants prevalent across the world. It is estimated that approximately 50% of UK dairy herds are infected with MAP, but this is likely an underestimate of the true prevalence. Infection can result in reduced milk yield, infertility and premature culling of the animal, leading to significant losses to the farming economy and negatively affecting animal welfare. Understanding the initial interaction between MAP and the host is critical to develop improved diagnostic tools and novel vaccines. Here we describe the characterisation of three different multicellular *in vitro* models derived from bovine intestinal tissue, and their use for the study of cellular interactions with MAP. In addition to the previously described basal-out 3D bovine enteroids, we have established viable 2D monolayers and 3D apical-out organoids. The apical-out enteroids differ from previously described bovine enteroids as the apical surface is exposed on the exterior surface of the 3D structure, enabling study of host-pathogen interactions at the epithelial surface without the need for microinjection. We have characterised the cell types present in each model system using RT-qPCR to detect predicted cell type-specific gene expression and confocal microscopy for cell type-specific protein expression. Each model contained the cells present in the bovine ileum and were therefore representative of the bovine gut. Exposure of the three model systems to the reference strain MAP K10, and a recent Scottish isolate referred to as C49, led to the observation of intracellular bacteria by confocal microscopy. Enumeration of the bacteria by genome copy number quantification, indicated that K10 was less invasive than C49 at early time points in infection in all model systems. This study shows that bovine enteroid-based models are permissive to infection with MAP and that these models may be useful in investigating early stages of MAP pathogenesis in a physiologically relevant *in vitro* system, whilst reducing the use of animals in scientific research.

## 1 Introduction

MAP is an acid-fast, Gram positive, facultative intracellular pathogen that causes Johne ‘s disease (JD) in ruminants. In cattle, symptoms will present 2-5 years after the initial infection and these include emaciation and chronic diarrhoea, which ultimately lead to death of the affected animal (Arsenault et al., 2014). This, coupled with reduced milk yield and fertility during the subclinical period, contributes to a significant burden on the farming economy. There are currently no effective vaccines to prevent MAP infection in cattle, and the available diagnostic tests routinely lack sensitivity and specificity. Understanding how MAP enters the host, and the mechanisms of the host-pathogen interaction are central to the development of new or improved methods for disease control.

MAP infection is most likely to occur in calves aged less than six months old through the ingestion of contaminated milk and colostrum (Bermudez et al., 2010; Windsor & Whittington, 2010). Horizontal transfer can also occur from the environment contaminated by other ruminants and wildlife reservoirs such as rabbits, foxes and stoats (Motiwala et al., 2004). Upon ingestion, MAP travels through the gastrointestinal tract to the small intestine where it can colonize the jejunum and ileum (Facciuolo et al., 2016). MAP has been shown to infect several cell types of the intestinal lining; including enterocytes (Pott et al., 2009), goblet cells (Schleig et al., 2005) and M cells (Bermudez et al., 2010; Ponnusamy et al., 2013), the latter of which have been hypothesised to be critical for infection of the host. Current dogma states that upon uptake by M cells, MAP can enter the underlying Peyer’s Patch of the small intestine where it is ingested by macrophages and dendritic cells (DCs) and survives and replicates in this niche (Bannantine & Bermudez, 2013; Khare et al., 2009). Cell death eventually leads to release of MAP and recruitment of additional macrophages and DCs to the tissue that also become infected (Arsenault et al., 2014; Köhler et al., 2015). The accumulation of infected immune cells drives a protective inflammatory response and the formation of granulomas in the intestinal lining (Tanaka et al., 2016). Onset of clinical disease results from an unknown trigger which causes a shift in host immune response from Th1 to Th2 dominant (Stabel et al., 2000), leading to an increase in bacterial shedding in the faeces and the onset of clinical signs.

Advances in the development of better diagnostic and identification of therapeutic targets have been limited by the lack of an appropriate model which is physiologically representative of a ruminant host, is reproducible and does not require, or minimises the use of, animals. The host species used to investigate MAP, the breed and age of animal, the route of infection, strain of MAP and inoculum dose all influence the outcome of experimental infections (Begg & Whittington, 2008). *In vivo* small animal models, including rabbits, mice and chickens, have been used for both short- or long-term infection studies (Kruiningen et al., 1991; Mokresh & Butler, 1990; Rosseeis et al., 2006). These models are more complex than monoculture cell lines and more easily housed than ruminants but are not necessarily representative of a natural MAP infection. These species require a higher or more prolonged dose of MAP to initiate an infection, and yet may not form granulomas in the intestine (Veazey et al., 1995). Therefore, an *in vitro* multicellular system representative of the bovine intestine is critical to advance MAP research. The generation of 3D bovine intestinal organoids (enteroids) has provided a model which maintains the diverse cell types present in bovine intestinal tissue whilst being cognizant of the principles of replacement, reduction and refinement (the 3Rs) underpinning more humane use of animals in UK research, due to their ability to be passaged (Hamilton et al., 2018b).

The aim of this study was to investigate if enteroid-derived models of the bovine intestine could be used to study the early host: pathogen events occurring following MAP infection. In addition to replicating the generation of 3D basal-out bovine enteroids (Hamilton et al., 2018), we describe the development of 2D monolayers and 3D apical-out bovine enteroids. These models were all representative of the intestinal cell lineages present in the animal tissue from which they were derived. Models were infected with the MAP K10 reference strain, and a recent cattle isolate from Scotland C49 (Mathie et al., 2020). Both strains successfully established an infection in all models, and whilst there was no evidence of bacterial replication over time, C49 consistently infected cells at higher levels than K10 suggesting greater infectivity or virulence. The models described here will provide an important system to dissect the host-pathogen interaction of the bovine gut with MAP.

## 2 Materials and Methods

### Animals

All tissues used in this study were obtained from healthy male British Holstein-Friesian (*Bos taurus*) calves (<1 year old). Calves were sourced from either the University of Edinburgh dairy herd, or from approved farms in Scotland. Ethical approval was granted by The Roslin Institute Animal Welfare Ethical Review Board.

### Generation and maintenance of 3D bovine enteroids

3D basal-out bovine enteroids were generated as previously described (Hamilton et al., 2018). Briefly, intestinal crypts were isolated from the terminal ileum of calves aged ≤9 months (n=5). Crypts were isolated by scraping the luminal side of the intestinal tissue. The crypts were subsequently washed in HBSS medium and digested at 37°C with DMEM medium containing 1% FBS, 25 μg/mL gentamicin, 20 μg/mL dispase I (Scientific Laboratory Supplies Ltd) and 75 U/mL collagenase (Merck Life Sciences) for 40 minutes, washed in HBSS and suspended in growth-factor reduced Matrigel (BD Biosciences, 356230). 50 μL of the suspension containing ∼200 crypts was plated in a 24-well tissue culture plate and was overlaid with 650 μL murine IntestiCult medium (STEMCELL Technologies) containing 10 μM each of a Rho-associated kinase inhibitor, Y27632 (Cambridge Bioscience), a p38 mitogen-activated protein kinase inhibitor SB202190 (Enzo Life Sciences) and a TGFβ inhibitor LY2157299 (Cambridge Bioscience) (henceforth referred to as complete IntestiCult medium). Enteroids were incubated at 37°C with 5% CO_2_ and passaged every 5-7 days of culture as previously described (Hamilton et al., 2018). 3D basal-out enteroids could be cryopreserved in Cryostor CS10 medium (STEMCELL Technologies) at -155°C for long-term storage. The enteroids were later resuscitated for experimental use as described previously (Hamilton et al., 2018).

### 2D monolayer generation

2D monolayers were established in glass 8 well-chambered slides coated in 2 μg/well bovine I collagen. Bovine intestinal crypts were isolated from bovine intestinal tissue (Hamilton et al., 2018), and were subsequently suspended in TrypLE Express (Thermo Fisher Scientific) and incubated at 37°C for 10 minutes. The crypts were disrupted using vigorous pipetting into a single cell suspension and were suspended in DMEM/F12 medium containing 1x B27 supplement minus vitamin A (Thermo Fisher Scientific) and 10% FBS. The cells were washed, re-suspended in complete IntestiCult medium and plated at a density of 2 × 10^5^ cells/well (Sutton et al., 2022). The cells were incubated at 37°C 5% CO_2_ and partial medium changes were performed at 48 hours post seeding, followed by full medium changes every 2-3 days thereafter.

### 3D apical-out enteroid generation

3D apical-out enteroids were established from previously passaged 3D basal-out enteroids and freshly isolated intestinal crypts. Passaged enteroids, or fresh crypts, were disrupted with TrypLE Express at 37°C for 10 minutes, washed and suspended in complete IntestiCult medium. The partially digested crypts were plated into wells of a 24-well plate at a density of ∼1000 clusters per well and incubated at 37°C with 5% CO_2_. After 24 hours, 3D apical-out enteroids were present in suspension, and the supernatant containing these enteroids were transferred a fresh well in a second 24 well plate. The medium was then partially changed every 2-3 days.

### Staining and imaging

Terminal ileum tissue was snap frozen in liquid nitrogen and embedded in OCT embedding matrix (CellPath UK Ltd). Seven micron thick slices were produced using a cryostat and adhered to microscope slides for fixation. Tissue slices, 3D enteroids and 2D monolayers were fixed with PBS containing 4% paraformaldehyde (PFA) for 1.5 hours at 4°C. Fixed cells were permeabilised in PBS containing 0.2% (v/v) Triton X-100 for 20 minutes and washed with PBS. Samples were then incubated in blocking buffer (PBS containing 0.5% bovine serum albumin (BSA) (v/v), 0.02% sodium azide (w/v) and 10% heat inactivated horse serum (v/v)) for 1 hour at room temperature. The samples were incubated with primary antibody (Supplementary Table S1) diluted in blocking buffer for 1 hour at room temperature, washed 3 times with PBS and incubated with the secondary antibody (Supplementary Table S1) diluted in blocking buffer for 1 hour at room temperature. Where specified, samples were stained with Phalloidin (Supplementary Table S2) diluted in blocking buffer. Samples were counterstained with DAPI for 5 minutes at room temperature, washed with distilled water, and mounted onto glass slides using ProLong Gold. The fluorescent signals were visualised using Leica LSM710 upright immunofluorescence microscope and images created in Zen Black software.

To stain mucins present in the enteroid models, samples were fixed in 4% PFA. The mucins were stained using Periodic Acid Schiff (PAS) staining according to the manufacturer’s instructions (TCS BioSciences Ltd), and counterstained with Scott’s tap water. The slide was then air-dried and mounted on a glass slide using Prolong Gold. Samples were imaged using a Brightfield microscope.

### RT-qPCR

Total RNA was extracted from all enteroid derived models, and intestinal tissue after the addition of a sterile steel ball and processing in the Qiagen Homogeniser for 3 minutes at 25 Hz, using Trizol (Thermo Fisher Scientific) according to the manufacturer’s instructions.

The cDNA was synthesised from the isolated RNA using Agilent AffinityScript Multiple temperature cDNA synthesis kit and confirmed to be free from contaminating genomic DNA using non-reverse transcriptase controls. The resulting cDNA synthesised from infected enteroid samples was diluted 1:20. Oligonucleotides were designed using Primer3 (Koressaar & Remm, 2007; Untergasser et al., 2012) and Netprimer (Biosoft International) software (see Supplementary Table S2 for primer sequences). The qPCR was performed using SYBR green Supermix (Quantabio, VWR International Ltd). The amplification of the primer sequence was performed at 60°C for 40 cycles and the generation of a single product was confirmed using dissociation curves.

The relative quantities of mRNA were calculated using the Pfaffl method (Pfaffl, 2001). A combination of reference genes were selected based on preliminary work to select genes with the smallest variation in gene expression between biological replicates and infection conditions. The RT-qPCR results for RALBP1 associated Eps domain containing 1 (REPS1) (Jensen et al., 2018) and actin beta (ACTB) were used as reference genes based on preliminary work demonstrating stable expression across infection conditions. The geometric mean of expression of these genes was used to calculate differences in the template RNA levels for standardisation of the Ct values for the genes of interest.

### Infection of bovine cells with MAP

MAP strains were cultured in 7H9 Middlebrook broth supplemented with 1 μg/mL Mycobactin J (ID Vet), 0.1% glycerol (v/v) and 0.1% Tween-80 (w/v). The bacteria were cultured at 37°C 100 rpm until an OD_600_ 0.6 was reached. 1 mL aliquots were frozen at -80°C until later use (Jensen et al., 2019; Mathie et al., 2020). For infection work, aliquots of MAP K10 or C49 were resuscitated from - 80°C storage by incubation at 37°C for 16 hours and then passed through a 30x gauge needle 10 times and diluted in cell growth media to the desired concentration.

For infection studies, cells were treated as follows:

**Basal-out 3D enteroids** were released from the Matrigel, and disrupted by pipetting to expose the apical surface of the cells. The disrupted multicellular structures were incubated with MAP at an MOI of 100 for one hour at 37°C to allow adherence and uptake into cells. The cells were washed twice in complete IntestiCult medium, one aliquot taken for gDNA extraction, and the rest plated in Matrigel-Intesticult for incubation at 37°C 5% CO_2_ for a further 24 and 72 hours. The **apical-out enteroids** were infected in a similar manner, except that they were not disrupted by pipetting at the beginning of the infection experiments and were plated in complete IntestiCult medium. **2D monolayers** were infected at an MOI of 10 for one hour at 37°C 5% CO_2_. Cells were washed twice in complete IntestiCult medium to remove any non-adherent bacteria and either processed for gDNA extraction, or incubated at 37°C 5% CO_2_ for a further 24 and 72 hours.

### Genomic DNA isolation and quantification

Genomic DNA was isolated from 3D enteroids, 2D monolayers and 3D apical-out enteroids infected with MAP, or non-infected controls, after the initial one hour infection time point and 24 and 72 hours post infection. In all instances the cells were treated with TrypLE express and incubated at 37°C for 10 minutes. Samples were suspended in 180 μL enzymatic lysis buffer containing 40 mg/mL lysozyme for 6 hours at 37°C before being digested with proteinase K at 56°C for 2.5 hours. The gDNA was then extracted using a Qiagen DNeasy Blood & Tissue Kit as described by the manufacturer’s instructions.

To enumerate the number of bovine cells and MAP bacteria present in the samples, qPCR was performed using SYBR green Supermix (Quantabio, VWR international Ltd). Primers were designed against the bovine Spastin gene and the MAP F57 sequence element (Supplementary Table S2) both of which are single copy genes. Templates with a known concentration of Spastin and F57 were used to generate the standard curves, from which the number of bovine and MAP cells could be extrapolated. Negative controls (no cDNA) were included to verify the absence of contamination.

### Statistical Analysis

Results are expressed as the mean of biological replicates ± standard deviation (SD). Statistical analysis was performed in GraphPad Prism using a 1-way or 2-way ANOVA as specified followed by the appropriate post hoc test for statistical significance.

## 3 Results

### Development and characterization of bovine enteroid models

Hamilton et al 2018 previously described a method for the isolation of bovine intestinal 3D basal-out organoids from ileal crypts of healthy male Holstein-Friesian calves. The enteroids could be passaged multiple times *in vitro*, and could reproducibly be resuscitated from cryo-preserved stocks. Here we have extended the work presented by Hamilton et al., 2018 by developing 2D monolayers and apical-out enteroids and subsequently using these, and the 3D basal-out enteroids, in infection studies.

Small intestinal crypts were taken from the terminal ileum of healthy male calves aged <9 months. The crypts were embedded in Matrigel domes and cultured with complete Intesticult medium for 7 days before passaging, as described in Hamilton et al., 2018. Within 24 hours of seeding, the crypts sealed over and formed enterospheres, and buds of developing organoids were observed by day 3 (Fig 1A-B). Over the course of 7 days the lumen filled with debris from the sloughing of dead cells from villus tips, mimicking the *in vivo* intestine (Fig 1C).

**Figure 1.**
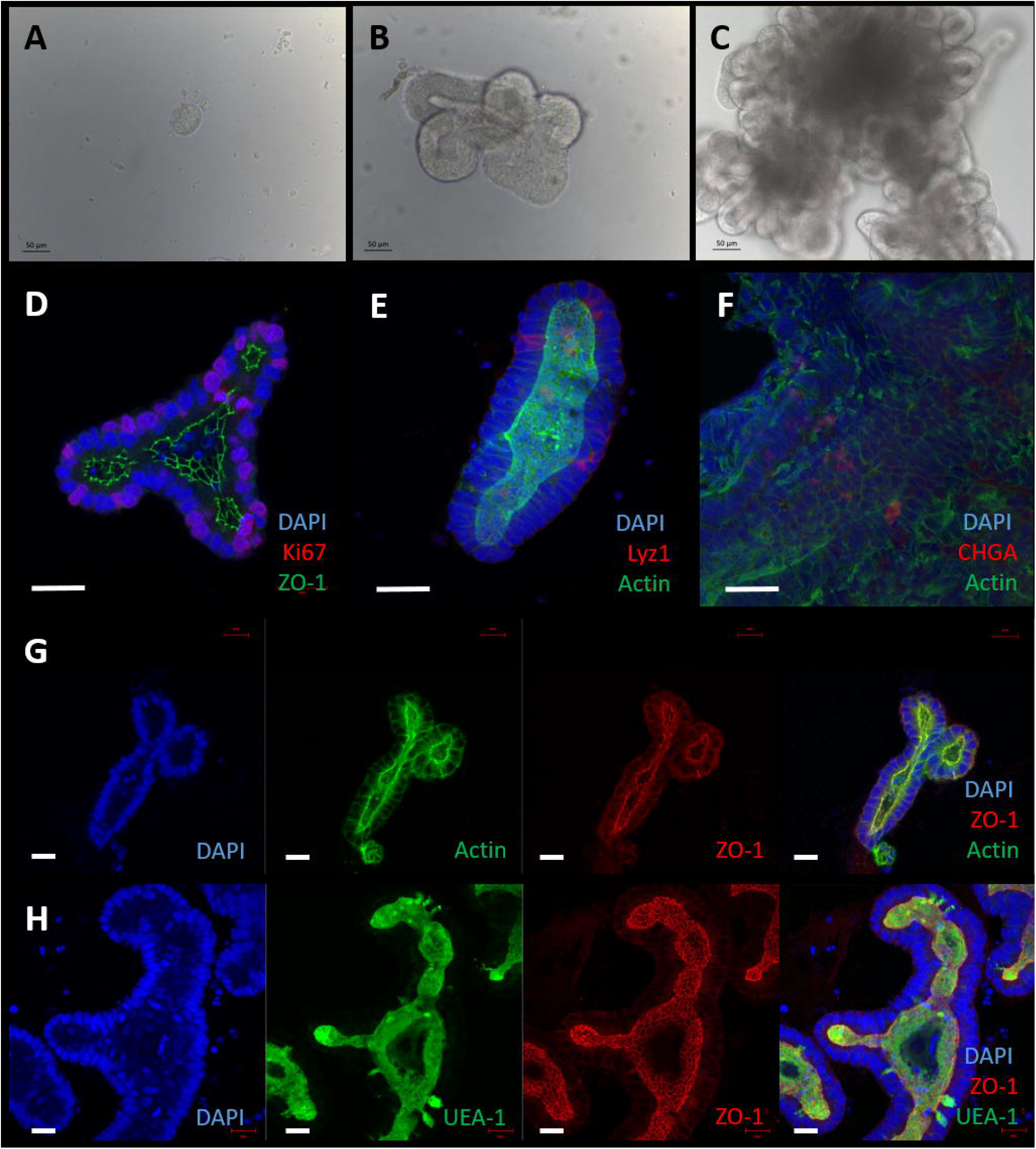
3D enteroids cultivated from bovine intestinal tissue. Representative images showing crypts isolated from bovine ileal tissue and maintained in a Matrigel dome with IntestiCult medium, from 5 independent animals. **(A-C)** Brightfield images of 3D enteroids cultured for 24 hours **(A)**, 3 days **(B)** and 7 days **(C)** demonstrating that they bud and proliferate over time. By 7 days the enteroid lumen fills with debris from cells sloughed off and are ready to be passaged. **(D-H)** Confocal images of 3D bovine enteroids stained for epithelial cell fate markers. **(D)** Nuclei (DAPI, blue), tight junctions (ZO-1, green) and proliferative cells (Ki-67, red). **(E)** Nuclei (DAPI, blue), actin (Phalloidin, green) and Paneth cells (lysozyme, red). **(F)** Nuclei (DAPI, blue), F-actin (Phalloidin, green) and enteroendocrine cells (chromogranin A, red). **(G)** Split panel of enteroids stained for nuclei (DAPI, blue), F-actin (Phalloidin, green), and tight junctions between cells (ZO-1, red). **(H)** Split panel of enteroids stained for nuclei (DAPI, blue), glycolipids (UEA-1, green) and tight junctions (ZO-1, red). Scale bar = 50 μm.

To ensure these basal-out enteroids contained the multiple cell lineages of the bovine intestine, predicted cell type-specific proteins were detected by immunofluorescence staining and confocal microscopy. Ki-67 (proliferative cell marker), lysozyme 1 (Paneth cell marker), chromogranin A (enteroendocrine cell marker), UEA-1 (lectin that binds glycoproteins and glycolipids with α-linked fucose residues) and ZO-1 (tight junctions) were confirmed to be present in the enteroids at 3 days of culture (Fig 1D-H). Positive F-actin staining with phalloidin was shown on the apical side of the cells facing the lumen (Fig 1 E-G), indicating the presence of a brush border and polarisation of the epithelium. The cell-type specific proteins observed in the enteroids using confocal microscopy reflects that observed in the original bovine intestinal tissue from which the organoids were derived (Supplementary Figure S1).

While enteroids offer a viable *in vitro* model of the bovine small intestine, there are limitations in their use for studies of MAP infection. In previously reported infection studies, enteroids have been microinjected directly into their luminal space (Bartfeld et al., 2015; Leslie et al., 2015; McCracken et al., 2014; Williamson et al., 2018). However, this is a time-consuming and laborious process, and is complicated by the irregular multi-lobular structures of bovine enteroids. With such a diverse range of enteroid size and shape, we could not infect them with a consistent number of bacteria per host cell (multiplicity of infection; MOI). To overcome this issue, we developed and characterised 2D monolayers and apical-out enteroids, which both offer a technically simpler way of interacting MAP with the apical surface of the cells without the need for microinjection.

To establish 2D monolayers, bovine intestinal crypts were enzymatically digested into a single cell suspension, and seeded onto collagen-coated wells of a tissue culture plate. The monolayers reached confluency by 3-4 days of culture and could be maintained for up to 10 days in complete Intesticult medium. To assess if these monolayers were representative of the cell lineages in bovine intestinal tissue, cell-type specific proteins were detected by immunofluorescence staining and confocal microscopy. Ki-67, lysozyme 1, chromogranin A, glycoproteins and glycolipids with α-linked fucose residues, F-actin and ZO-1 were confirmed to be present in the monolayers (Fig 2A-F). Periodic Acid Schiff staining was used to stain mucins characteristic of goblet cells (Fig 2D). These characteristics are consistent with the original bovine tissue from which they were derived (Supplementary Figure S1).

**Figure 2.**
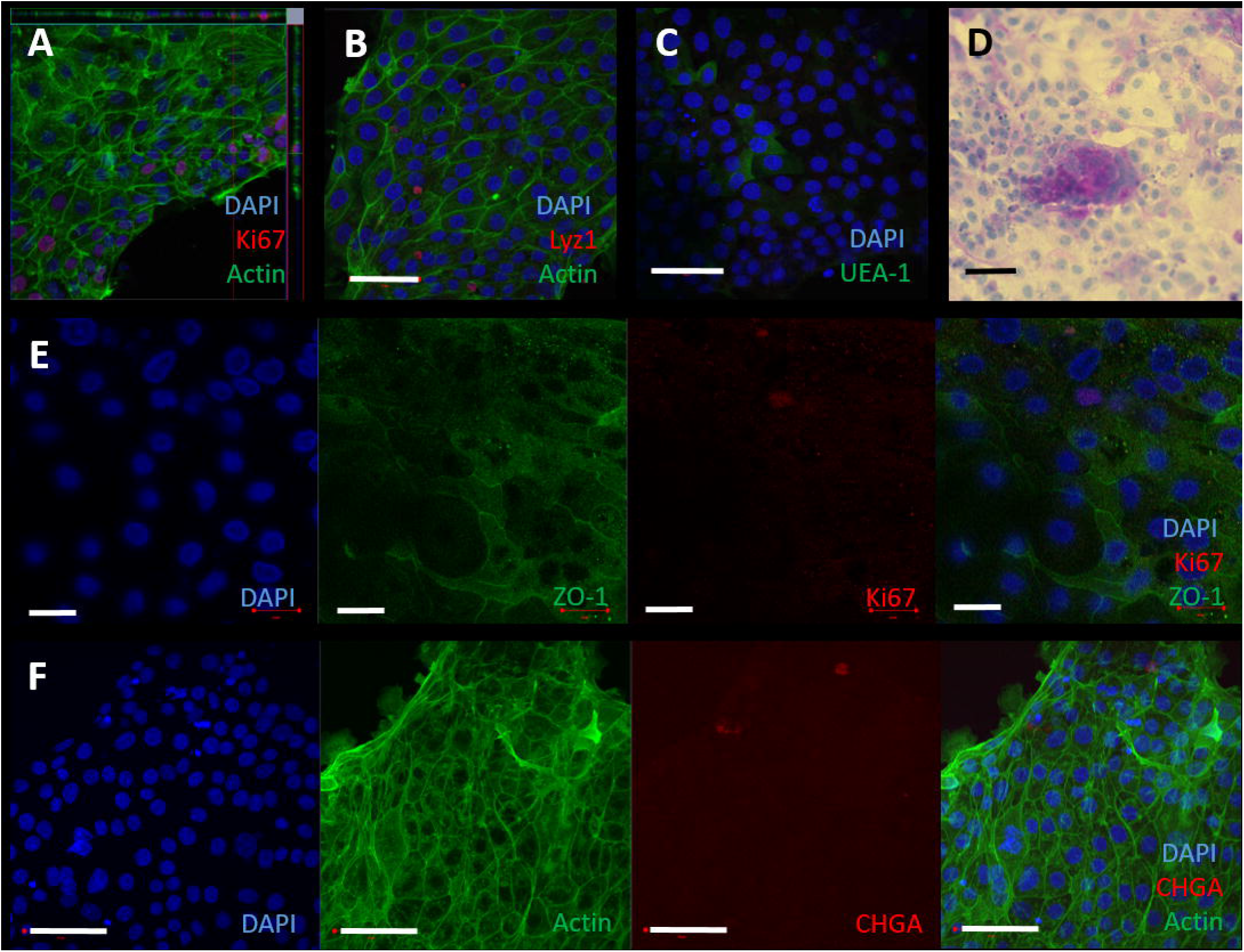
2D epithelial monolayers cultured on collagen matrix. Single cells were seeded onto collagen coated wells. The 2D monolayers were cultured with IntestiCult containing the relevant inhibitors and maintained for up to 10 days. **(A-F)** Representative confocal microscopy images of 2D monolayers cultured on collagen coated wells from 3 separate amimals, demonstrating presence of specific cell marker proteins. **(A)** F-actin (phalloidin, green), nuclei (DAPI, blue) and proliferative cells (Ki67, red). **(B)** Nuclei (DAPI, blue), F-actin (Phalloidin, green) and Paneth cells (lysozyme, red). **(C)** Nuclei (DAPI, blue) and glycolipids (UEA-1, green). **(D)** 2D monolayer imaged by brightfield microscopy stained for Periodic Acid Schiff to show mucins produced by goblet cells. **(E)** Split panel of monolayer stained for nuclei (DAPI, blue), tight junctions between cells (ZO-1, green), and proliferative cells (Ki-67, red). Scale bar = 20 μm. **(F)** Split panel of monolayer stained for nuclei (DAPI, blue), F-actin (Phalloidin, green) and enteroendocrine cells (chromogranin A, red). Scale bar = 50 μm.

When we observed the 2D monolayers over time, we noted the presence of 3D structures in the medium overlay. These displayed a characteristically different morphology to 3D enteroids suspended in Matrigel (Fig 3A-B). It was hypothesised these may be apical-out enteroids due to their morphology and formation in suspension, which is characteristic of apical-out avian enteroids described recently (Nash et al., 2021). The structures in suspension were impermeable to 4 kDa FITC-dextran in the absence of EDTA (Fig 3C-D), indicating the presence of intact cell: cell tight junctions. To confirm the apical-out polarisation, the bovine enteroids were stained with Phalloidin to localise F-actin (Fig 3E). F-actin expression was observed on the outer surfaces of these enteroids, in contrast to basal-out 3D enteroids cultured in Matrigel where F-actin was on the luminal side of the structure (Fig 1E).

**Figure 3.**
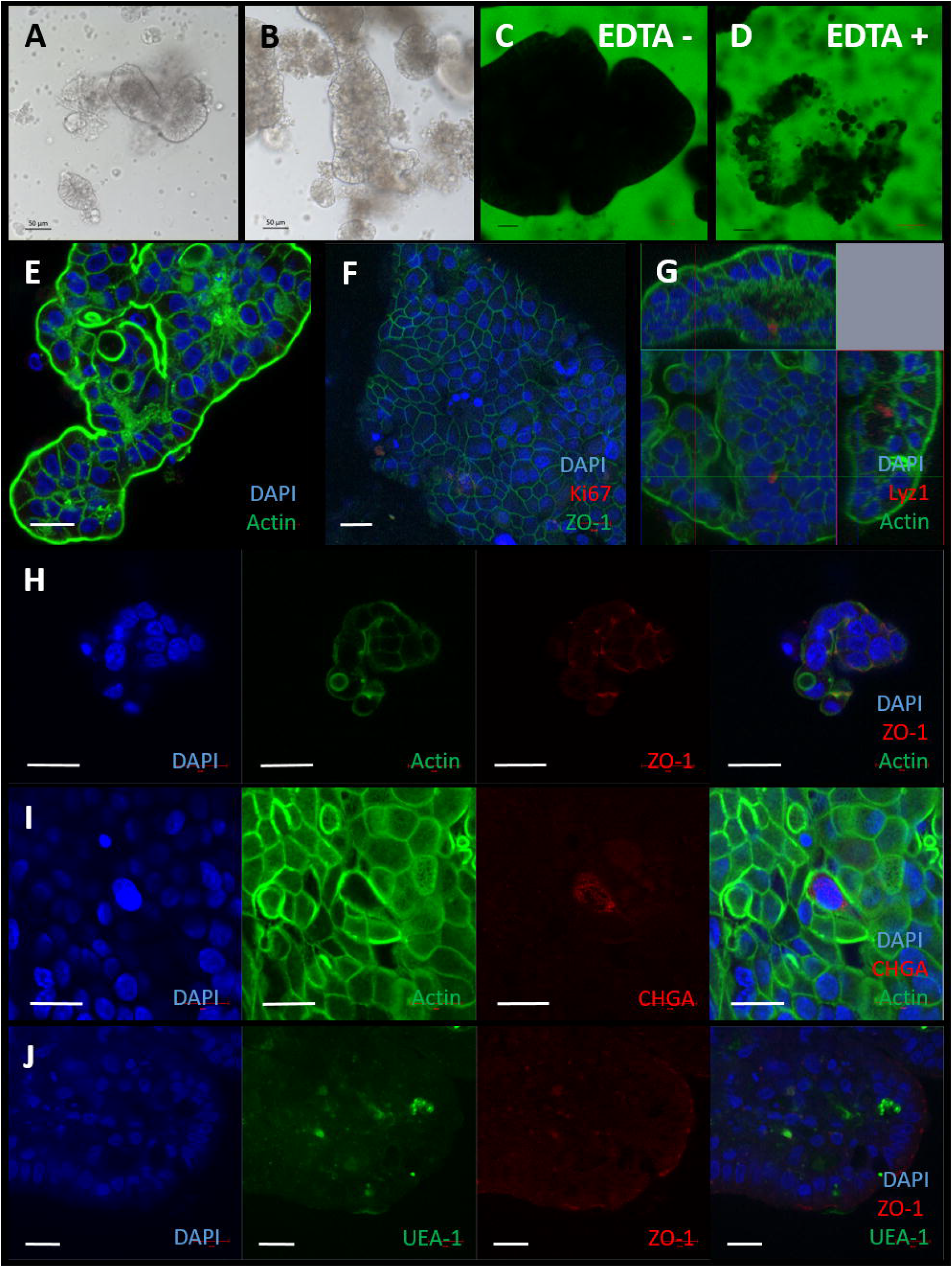
Apical-out bovine enteroids show epithelial barrier integrity when established from freshly harvested intestinal crypts. Apical-out enteroids cultivated from freshly isolated intestinal crypts and cultured in suspension in Intesticult at 1 day **(A)** and 7 days **(B)**. Images are representative of enteroids derived from 3 independent animals. **(C-D)** Confocal images of bovine inside out enteroids (7 days of culture) immersed in FITC-dextran 4kDa showing epithelial barrier integrity in untreated **(C)** and EDTA-treated conditions **(D)**. Immunofluorescence staining of bovine inside out intestinal organoids shown in split panel demonstrates epithelial differentiation **(E-J)**. Apical-out enteroids show reverse polarisation compared to basal-out 3D enteroids from the brush border facing the external medium, represented by F-actin staining (Phalloidin, green); and a dense internal core of cells (nuclei stained with DAPI, blue) **(E)**. Nuclei (DAPI, blue), proliferative cells (Ki67, red), and tight junctions between cells (ZO-1, green) **(F)**. Cross section of an apical-out enteroid from z-stack images of apical-out enteroids stained for nuclei (DAPI, blue), F-actin (Phalloidin, green) and Paneth cells (Lysozyme 1, red) **(G)**. Split panel of apical-out enteroids stained for nuclei (DAPI, blue), actin (Phalloidin, green) and tight junctions between cells (ZO-1, red) **(H)**. Split panel of apical-out enteroids stained for nuclei (DAPI, blue), F-actin (Phalloidin, green) and enteroendocrine cells (Chromogranin A, red) **(I)**. Split panel of apical-out enteroids stained for nuclei (DAPI, blue), glycoproteins/glycolipids (UEA-1, green) and tight junctions (ZO-1, red) **(J)**. Scale bar = 20 μm.

Antibody staining and confocal microscopy was used to demonstrate the presence of cell-specific proteins of apical-out enteroids at 7 days of culture. Lysozyme 1, chromogranin A, glycoproteins and glycolipids with α-linked fucose residues and ZO-1 were present (Fig 3G-J). One major difference between the 3D apical-out enteroids and the other two cell models described in this paper, was a lack of Ki-67 staining by 7 days of culture (Fig 3F).

Apical-out enteroids were also generated from previously passaged enteroids by culture in suspension. These apical-out enteroids were more spheroid in appearance than those generated from fresh crypts (compare Fig 3 with Supplementary Figure S2A-B), but were still impermeable to 4kDa FITC-dextran (Fig S2C-D). After 7 days culture the apical-out enteroids were antibody stained and imaged by confocal microscopy. All of the expected mature epithelial cell types were identified (Fig SE-H). However, the data demonstrated that this culture method generated a mixed culture of basal-out and apical-out 3D enteroids (Supplementary Figure S2I). For this reason, apical-out 3D enteroids generated from fresh harvested intestinal crypts were used for MAP infection studies.

Studies of livestock species is inherently limited by the lack of specific reagents, particularly antibodies for cell type-specific molecules. This has limited the capability to determine the precise nature of (for example) bovine M cells. Therefore, to extend the data presented in Figures 1 to 3, we utilised qRT-PCR to detect expression of genes predicted to be specific to certain cell types, and to compare their abundance between the three models described herein (Fig 4). As a definitive mRNA or cell surface expression signature profile for bovine M cells remains unknown, three different genes which have been reported in the literature as specific to M cells were investigated. GP2 is a known murine M cell transcript (Ohno & Hase, 2010), whilst cyclophilin A (PPIA) and cytokeratin 18 (KRT18) are both predicted bovine M cell transcripts (Hondo et al., 2011, 2016). In other species’ enteroid models, M cells are absent from the system until they are stimulated with recombinant RANK-L. All three gene transcripts were significantly upregulated in the basal-out 3D enteroids compared to the tissue of origin, particularly KRT18. Given the current literature, and the fact that we have been unable to demonstrate the presence of phagocytic cells in our systems, we do not believe these are M cell specific gene transcripts. Interestingly, 2D monolayers did not show an upregulation in expression of PPIA or GP2.

**Figure 4.**
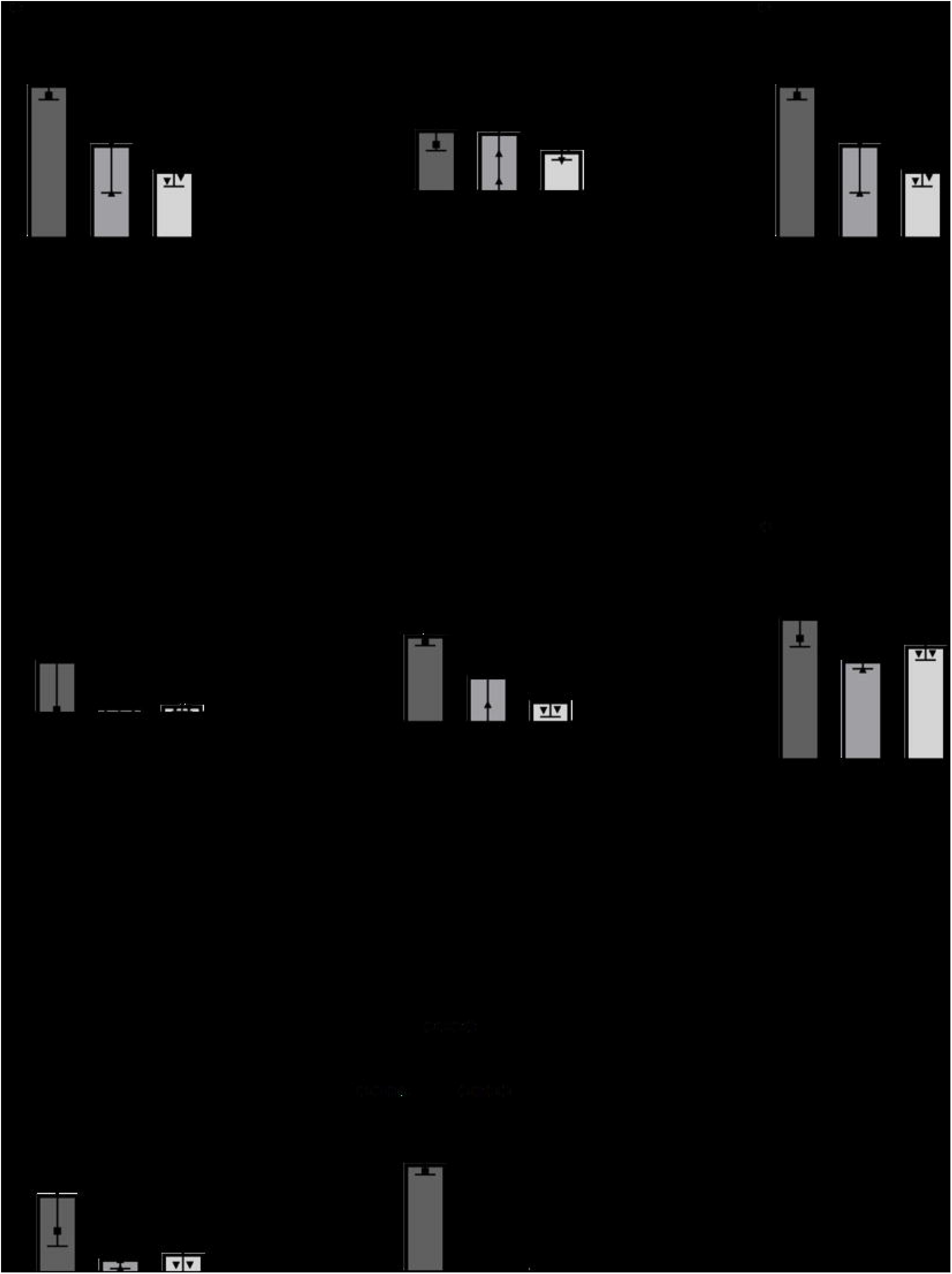
RT-qPCR of mRNA expression indicative of specific cell types in enteroid-derived models. The gene expression was determined by RT-qPCR and calculated as fold change relative to the expression of *ACTB* and *REPS1* as endogenous reference genes. Total RNA was isolated from the samples and confirmed to be free of genomic DNA. The results are from samples derived from 2 independent animals, with 3 technical repeats and presented as mean values ± standard deviation (SD). Statistical analysis was performed with a one-way ANOVA; * = P≤0.05; **=P≤0.01; ***=P≤0.001; ****=P≤0.0001.

For the genes indicative of enterocytes (*VIL1*), stem cells (*LGR5*), Paneth cells (*LYZ1*), goblet cells (*MUC2*) and enteroendocrine cells (*CHGA*), transcripts were detected in both the original bovine intestinal tissue samples and the enteroid models (Fig 4), although significant differences in expression levels between tissue and specific enteroid models was also observed.

The apical-out enteroids and the 2D monolayer demonstrated a decreased abundance of LGR5+ stem cells compared to the 3D basal-out enteroids and intestinal tissue. This corresponds with a reduced abundance of LYZ1+ paneth cells which help support the stem cell niche. This is not unexpected as these models showed a reduced capacity to proliferate for long periods of time. Interestingly, an upregulation of CHGA gene expression in 3D enteroids indicated an increased abundance of enteroendocrine cells, which may reflect specific culture conditions that support the maintenance of this particular cell type. However, this abundance was not observed at the protein level upon staining for CHGA in 3D basal-out enteroids (Fig 1F), and therefore it is likely that while the upregulation of CHGA gene expression is significant, there was no increase in the number of enteroendocrine cells in the model.

### MAP infection of enteroid models

To investigate whether the three enteroid-derived models were permissive to MAP infection, the MAP K10 reference strain was compared with C49, a recent Scottish cattle MAP strain isolated from a local farm (Mathie et al., 2020). Due to the prolonged length of time required to quantify viable MAP in culture, the total number of bacteria was measured using qPCR of the F57 sequence element from genomic DNA to determine the genome copy number. Similarly, the bovine cell number was measured by qPCR of the spastin gene from genomic DNA to determine the genome copy number.

3D basal-out enteroids were infected at the same time as they were passaged with identical numbers of viable bacteria for each strain of MAP. By 72 hours, the number of bovine cells had significantly increased from 0 and 24 hours post-infection in the non-infected and MAP K10 infected conditions, while there was a trend of fewer bovine cells being present in samples infected with MAP C49 (Fig 5A). At 24 hours, there were significantly higher numbers of MAP C49 present compared to MAP K10 (Fig 5B). By 72 hours this difference between strains remained evident, but was no longer statistically significant.

**Figure 5.**
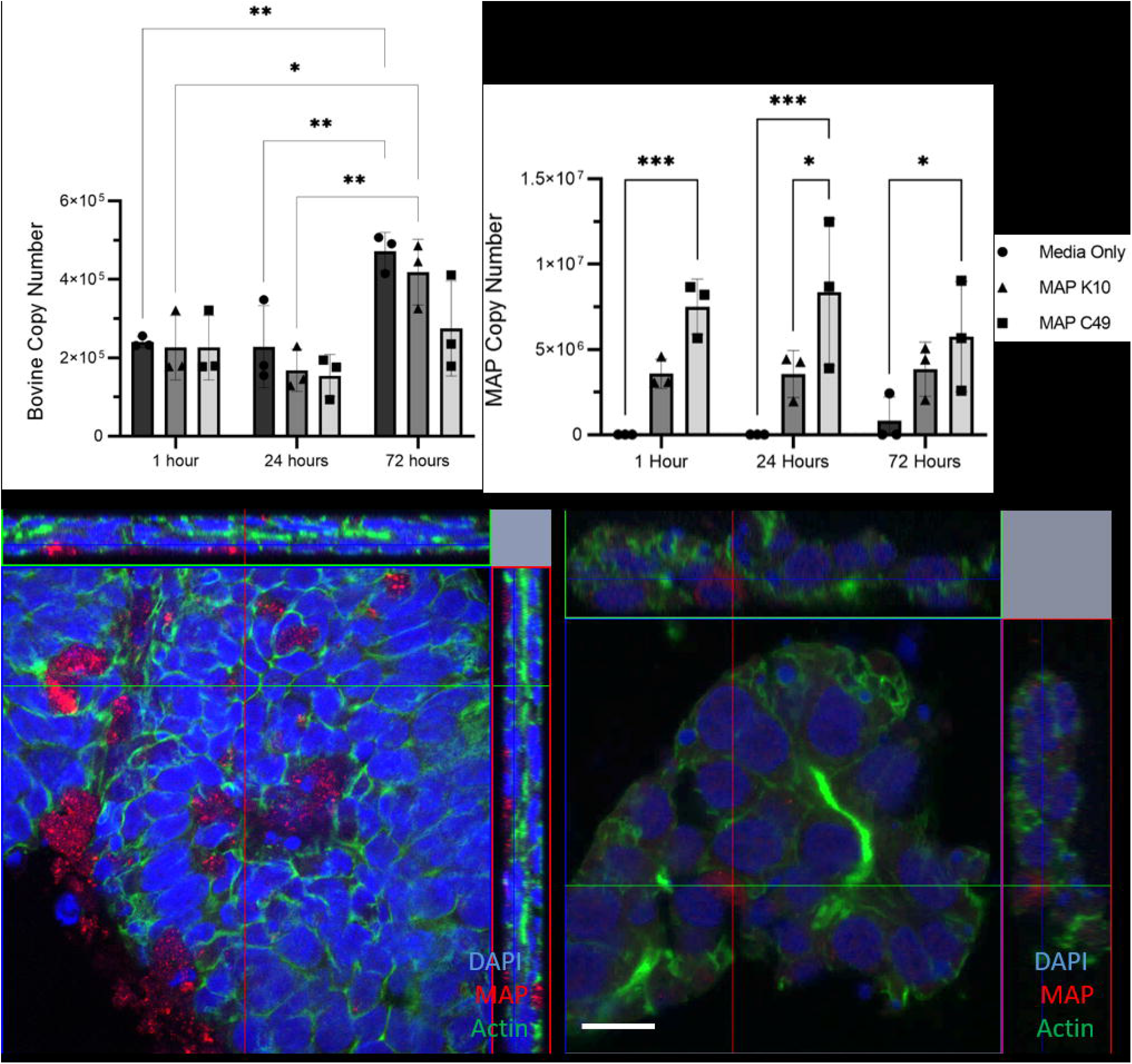
Infection of basal-out 3D enteroids with 2 different strains of MAP. Enteroids were infected with an MOI of 100 in 3 biological replicates using enteroids generated from the same calf at passage 15, 17 and 20. The bovine **(A)** and bacterial **(B)** cell number was quantified using qPCR for the spastin gene and F57 sequence element respectively via qPCR (data presented as mean ± SD). Statistical analysis performed using 2-way ANOVA followed by a post hoc Tukey’s test. P<0.05 = *; P<0.01 = **; P<0.001 = ***; P<0.0001 = ****. **(C-D)** Immunofluorescence microscopy images of infected 3D enteroids 24 hours post infection. Enteroids were stained for nuclei (DAPI, blue), F-actin (Phalloidin, green) and MAP (anti-MAP, red). 3D enteroids are shown as a cross section from Z-stack images infected with MAP K10 **(C)** and MAP C49 **(D)**. Scale bar = 10μm.

At 24 hours post infection, enteroids were stained and analysed by immunofluorescence confocal microscopy. Both MAP K10 and C49 were shown to be intracellular at this time indicating this model is permissive to infection (Fig 5C-D). However, many bacteria were also extracellular, trapped in the lumen of the enteroid structure or adhered to the basolateral side of the enteroids.

In this model, MAP C49 remained present in the bovine enteroids in greater numbers than MAP K10 over the course of the experiment. This may indicate that the initial infection for only 1 hour at the beginning of the time course was more effective for the C49 strain than K10. Over the course of infection, the numbers of K10 and C49 do not significantly change, although there is a small decrease in C49 from 24 hours to 72 hours post-infection (Fig 5B). This data could indicate that there is no evidence of bacterial replication in these cultures, and there is also little evidence of bacterial killing by cell-autonomous mechanisms such as autophagy or pyroptosis.

2D monolayers were infected with identical numbers of viable MAP inoculated directly into the cell medium for 1 hour at 37°C, before being washed three times with PBS to remove non-adherent bacteria. Genomic DNA samples were extracted at this point (1 hour), and replicate samples incubated for a total of 24 and 72 hours prior to DNA extraction. Upon quantifying the host and bacterial genome copy numbers by qPCR, a similar trend to the 3D enteroid model was observed (Fig 6 A-B). Significantly more MAP C49 bacteria were present at 1 hour post-infection compared to MAP K10 (Fig 6B). At 24 and 72 hours post infection this difference remained but was no longer statistically significant. Interestingly, the bacterial cell numbers increase over time for both MAP K10 and MAP C49, indicating replication of the bacteria in this model, but this was not statistically significant. The number of bovine cells remained constant over 72 hours, consistent with a lack of proliferative capacity upon reaching confluency. Similar to the data shown in Fig 5, intracellular bacteria could be observed by immunofluorescence microscopy, with some bacteria in an extracellular compartment likely attached to the outer surface of the cells or directly to the collagen matrix (Fig 6C-D).

**Figure 6.**
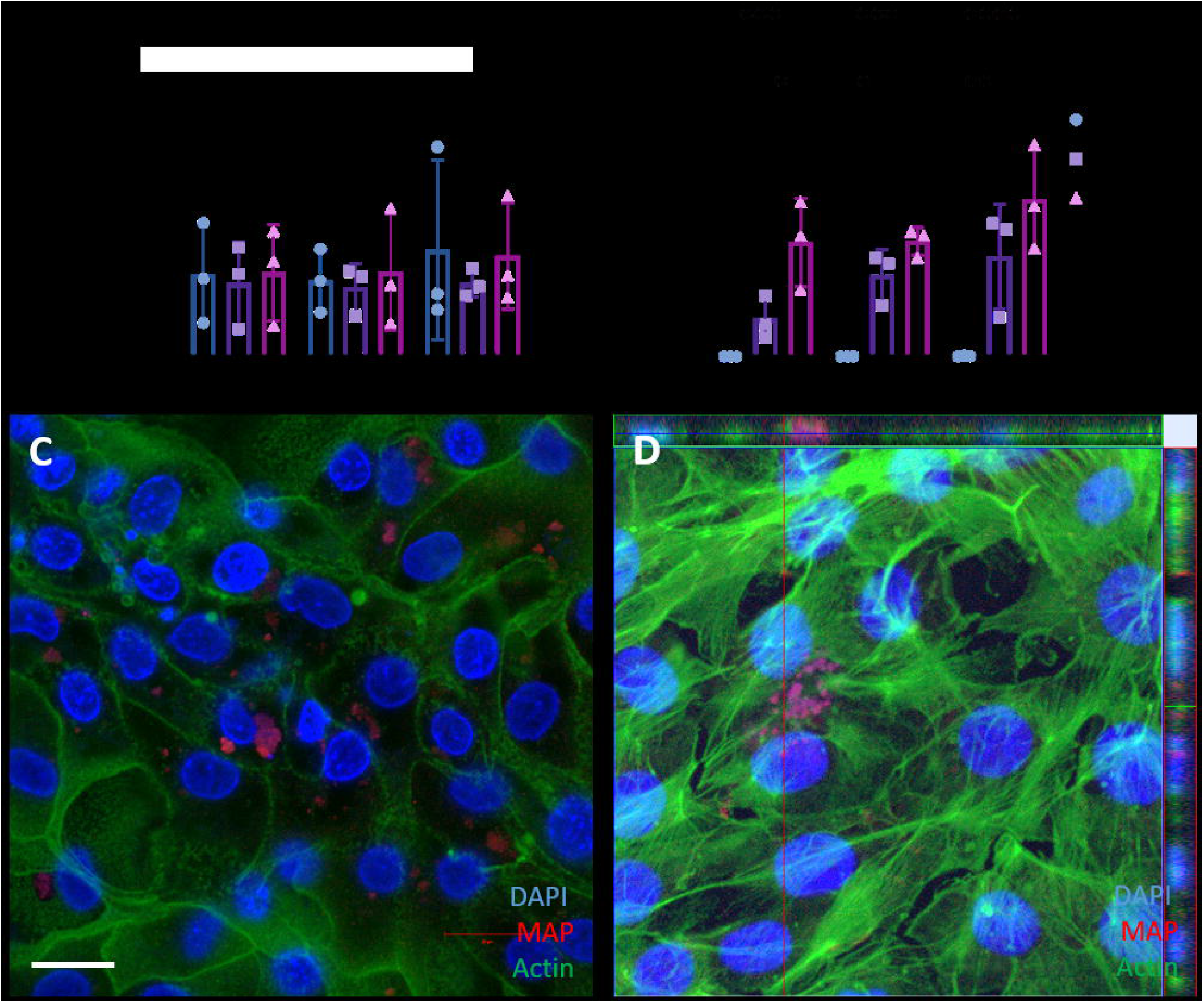
Infection of 2D monolayers with 2 different strains of MAP. Monolayers were infected with an MOI of 10 in 3 independent experiments. The bovine **(A)** and bacterial **(B)** cell number was quantified using qPCR for the spastin gene and F57 sequence element respectively via qPCR (data presented as mean ± SD). Statistical analysis performed using 2-way ANOVA followed by a post hoc Tukey’s test. P<0.05 = *; P<0.01 = **; P<0.001 = ***; P<0.0001 = ****. **(C-D)** Immunofluorescence staining of monolayers imaged 24 hours post infection. Infected monolayers were fixed and stained for nuclei (DAPI, blue), F-actin (Phalloidin, green) and MAP (anti-MAP, red). Cells were infected with MAP K10 **(C)** and as a cross section from Z-stack images infected with MAP C49 **(D)**. Scale bar = 10μm.

Similarly, 3D apical-out enteroids were infected with MAP by inoculating the culture medium with identical numbers of the two MAP strains. Over the course of 72 hours the bovine cell number remained constant, as expected for a model lacking proliferative cells (Fig 7A). As with the other bovine enteroid models, significantly higher numbers of MAP C49 were present at 1 hour and 24 hours post-infection when compared with MAP K10 (Fig 7B). Despite demonstrating that the bacteria were intracellular by 24 hours post-infection (Fig 7C-D), there was no significant increase in bacterial numbers over time, indicating that the bacteria are unlikely to be actively replicating in this model.

**Figure 7.**
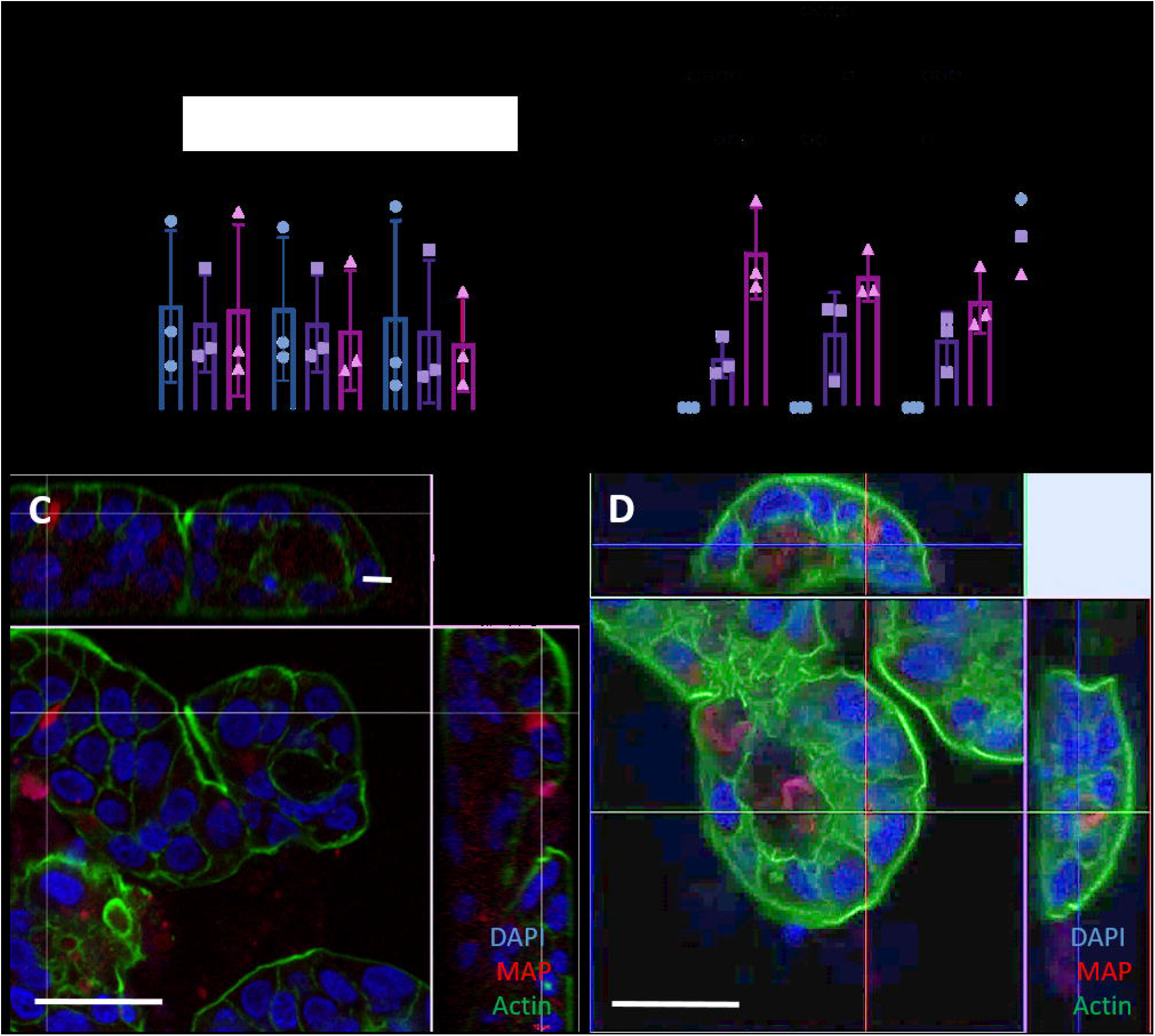
Infection of 3D Apical-out enteroids with 2 different strains of MAP. Apical-out enteroids were infected with an MOI of 100 using apical-out enteroids generated from 3 separate calves. The bovine **(A)** and bacterial **(B)** cell number was quantified using qPCR for the spastin gene and F57 sequence element respectively via qPCR (data presented as mean ± SD). Statistical analysis performed using 2-way ANOVA followed by a post hoc Tukey’s test. P<0.05 = *; P<0.01 = **; P<0.001 = ***; P<0.0001 = ****. **(C-D)** Immunofluorescence staining of 3D apical-out enteroids imaged 24 hours post infection. Slides with infected apical-out enteroids were fixed and stained for nuclei (DAPI, blue), F-actin (Phalloidin, green) and MAP (anti-MAP, red). Apical-out enteroids were infected with MAP K10 **(C)** and as a cross section from Z-stack images infected with MAP C49 **(D)**. Scale bar = 10μm.

## 4 Discussion

Here we describe the establishment and culture of bovine 2D monolayers from bovine small intestinal crypts. We also demonstrate the formation of 3D apical-out enteroids directly from bovine intestinal crypts or previously passaged 3D basal-out enteroids. These multi-cellular cultures were characterised by immunofluorescence staining and confocal microscopy (where bovine-reactive reagents exist) and RT-qPCR for predicted cell lineage markers, to identify the presence of multiple intestinal cell types. With the exception of proliferative cells, all three model systems maintained a similar cellular composition to the bovine intestinal tissues from which they were derived.

The primary aim of the work presented here was to develop a model system to interrogate the interaction of MAP with cells of the bovine intestine. Given the technical difficulties associated with microinjection of 3D organoids, we have shown that 2D monolayers and apical-out enteroids would be useful in this regard. Interestingly, irrespective of the type of enteroid-derived model system we infected, our recent cattle MAP isolate C49 was capable of higher levels of initial infection of the cells, when compared to K10. This could reflect the finding that K10 is laboratory adapted to a point of significantly lower virulence (Radosevich et al., 2006). By quantifying bacterial and host cell genomes, we were able to gain valuable insight into the fate of both bacteria and bovine cells over time. No evidence of significant bacterial replication, which would be indicated by increasing numbers of bacteria over time, was detected. Similarly bovine cell numbers were maintained (and even increased over time in the basal-out 3D cultures), with no evidence of induction of inflammatory cell death pathways such as pyroptosis that are often associated with control of intracellular pathogens. A significant caveat to this method is that we cannot assume that every MAP genome that we quantified from the cell cultures originates from a viable bacterium. There are several advantages and disadvantages of the different methods of MAP quantification used by different groups around the world. In addition to the length of incubation time to obtain colonies, bacterial culture can lead to an underestimation of the number of viable bacteria due to the propensity of MAP to aggregate. Methods based on confocal microscopy (Mathie et al., 2020), or genome copy number, may not differentiate between live and dead bacteria.

Unfortunately, the limited availability of reagents for bovine cell markers has hindered the identification of certain cell-specific molecules in the bovine system such as M cells. As an important cell targeted by multiple enteric pathogens such as *Salmonella enterica*, reliable stimulation with bovine RANK-L treatment and unequivocal identification of M cells in samples, would represent significant progress to our understanding of multiple bovine intestinal infections. Furthermore, reagents specific to a wide range of bovine cell types would allow the study of cell tropism displayed by these pathogens in a physiologically representative system. Recent work has described the incorporation of a microbiome (Puschhof et al., 2021), in addition to co-cultures with immune cells (Takashima et al., 2019), with some researchers pursuing replicating the intraluminal “flow” and peristaltic movement of an intestine with “organ-on-a-chip” system (Kasendra et al., 2018; Kim et al., 2012).

Overall, the establishment of these bovine intestinal cultures has provided novel, physiologically relevant *in vitro* models that offer multiple routes of infection depending on the researcher’s preference. Our data implies that these enteroids cultures are amenable to the study of many enteric pathogens, including MAP, providing crucial understanding of host-pathogen interactions in the bovine small intestine. Finally, these models offer attractive solution to bridge basic and translational research, whilst importantly reducing the use of animals in research.

## Figure Legends

**Supplementary Figure 1.**
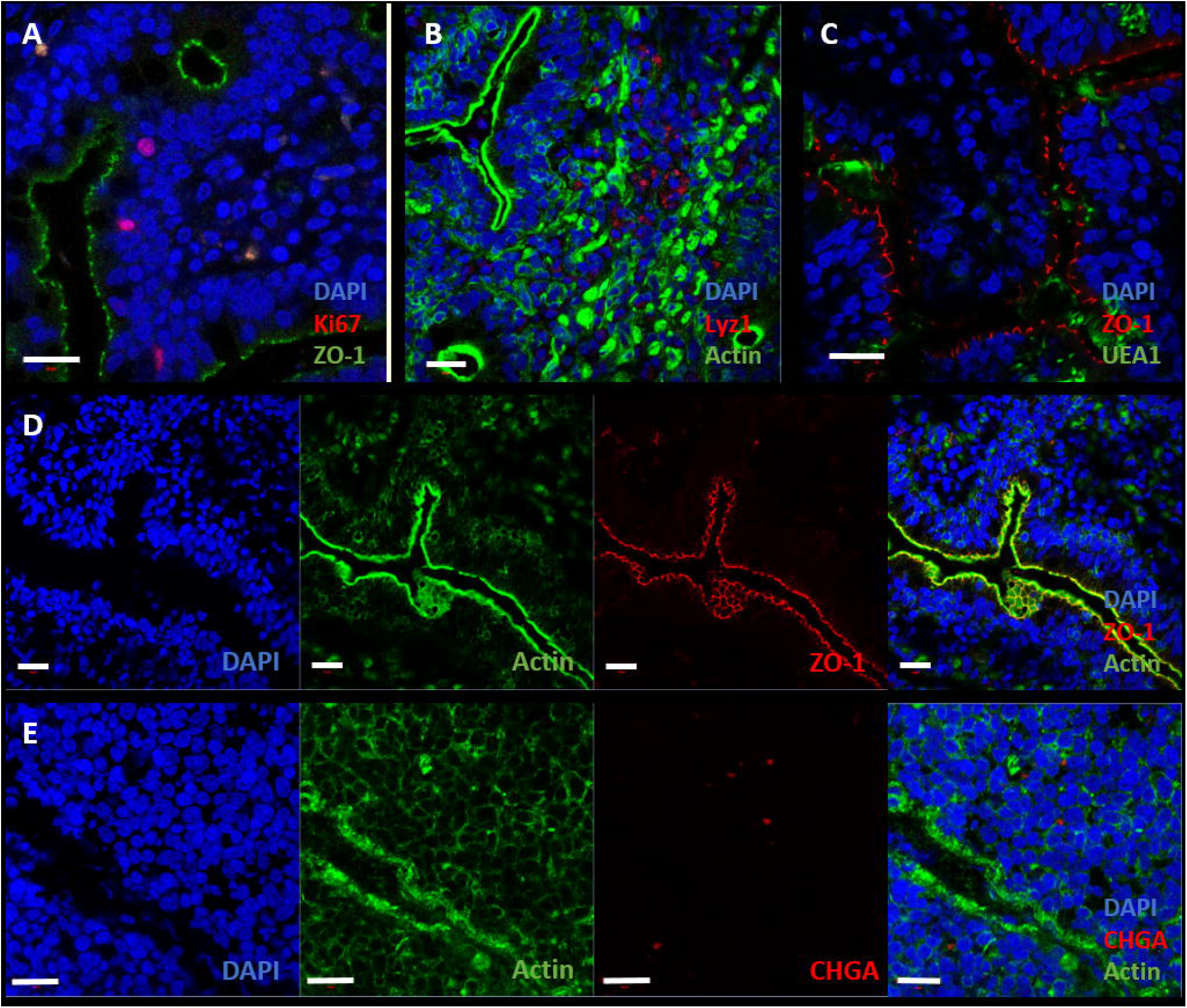
Immunofluorescence staining of bovine intestinal tissue slices. Representative confocal images from 2 calves showing staining for epithelial cell fate markers. **(A)** Tissue slices stained for nuclei (DAPI, blue), tight junctions (ZO-1, green) and proliferative cells (Ki-67, red). **(B)** Tissue stained for nuclei (DAPI, blue), actin (Phalloidin, green) and Paneth cells (lysozyme, red). **(C)** Tissue stained for nuclei (DAPI, blue), tight junctions (ZO-1, red) and glycolipids (UEA-1, green). **(D)** Split panel of tissue stained for nuclei (DAPI, blue), actin (Phalloidin, green), and tight junctions between cells (ZO-1, red). **(E)** Split panel of tissue stained for nuclei (DAPI, blue), acting (phalloidin, green) and enteroendocrine cells (chromogranin A, red). Scale bar = 20 μm.

**Supplementary Figure 2.**
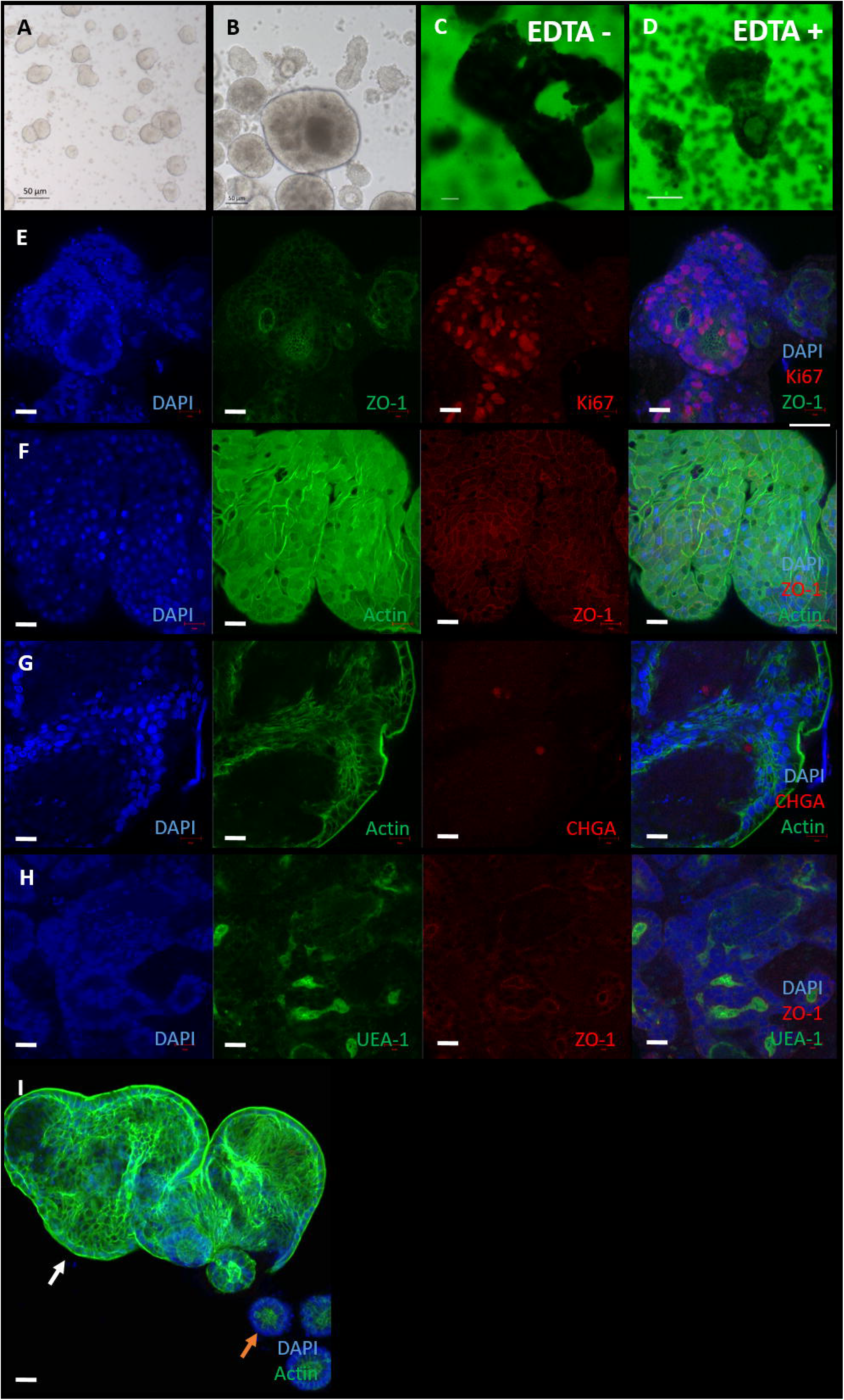
Apical-out bovine enteroids show epithelial barrier integrity when established from previously passaged 3D enteroids. Apical-out enteroids established from previously passaged basal-out 3D enteroids at 1 day **(A)** and 7 days **(B)**. Images are representative of enteroids generated from 1 calf at passage number 5, 11 and 13. **(C-D)** Confocal images of bovine apical-out enteroids (7 days of culture) immersed in FITC-dextran 4kDa showing epithelial barrier integrity in untreated **(C)** and EDTA-treatment **(D)**. Scale bar = 50 μm. Immunofluorescence staining of bovine apical-out enteroids shown in split panel demonstrates epithelial differentiation **(E-H)**. Apical-out enteroids were stained for nuclei (DAPI, blue), proliferative cells (Ki67, red), and tight junctions between cells (ZO-1, green) **(E)**. Apical-out enteroids were stained for nuclei (DAPI, blue), tight junctions (ZO-1, red), and actin (Phalloidin, green) **(F)**. Apical-out enteroids stained for nuclei (DAPI, blue), actin (Phalloidin, green) and enteroendocrine cells (Chromogranin A, red) **(G)**. Apical-out enteroids stained for nuclei (DAPI, blue), glycolipids (UEA-1, green) and tight junctions (ZO-1, red) **(H)**. Apical-out enteroids and basal-out 3D enteroids shown to be present in the same culture when generated from previously passaged enteroids (Phalloidin, green and DAPI, blue) **(I)**. White arrow denotes apical-out enteroid, orange arrow denotes basal-out enteroid. Scale bar = 20 μm.

## 6 Author Contributions

All authors agree to be accountable for all aspects of the work in ensuring that questions related to the accuracy or integrity of any part of the work are appropriately investigated and resolved. Conceptualization: RB and JS. Investigation, Project administration and Writing - original draft: RB and JS. Formal analysis, Validation and Visualization: RB and KJ. Methodology: RB and KJ. Funding acquisition, Resources, Supervision: JS, NM and JH. Writing - review & editing: JS, KJ, and JH. All authors contributed to the article and approved the submitted version.

## 8 Acknowledgments

We would like to acknowledge allege our colleagues for their comments and suggestions for experimental design. In particular, we are grateful to Dr. Rachel Young for her contributions of time and expertise.

## 9 Supplementary Material

Supplementary Material should be uploaded separately on submission, if there are Supplementary Figures, please include the caption in the same file as the figure. Supplementary Material templates can be found in the Frontiers Word Templates file. Please see the [Supplementary Material section of the Author guidelines](https://www.frontiersin.org/about/author-guidelines#SupplementaryMaterial) for details on the different file types accepted.

## 1 Data Availability Statement

The datasets [GENERATED/ANALYZED] for this study can be found in the [NAME OF REPOSITORY] [LINK]. Please see the [Data Availability section of the Author guidelines](https://www.frontiersin.org/about/author-guidelines#AvailabilityofData) for more 657 details.

---

**Supplementary Table S1.** Table of cell staining reagents for confocal microscopy.

| **Antibody/ cell stain** | **Concentration for use in confocal microscopy** | **Supplier name and product code** |
| --- | --- | --- |
| Anti-Ki-67 | 10 µg/mL | Abcam, ab15580 |
| Anti-ZO-1 | 5 µg/mL | Thermo Fisher, 1A12 |
| Anti-Lysozyme 1 | 10 µg/mL | Dako, A0099 |
| Anti-Chromogranin A | 20 µg/mL | Santa Cruz biologicals, SC-271738 |
| *Ulex Europaeus* Agglutinin 1 | 10 µg/mL | Vector Laboratories, FL-1061 |
| Anti-MAP | 3 µg/mL | Gene Tex, GTX82978 |
| Phalloidin 488 | 66 µM | Thermo Fisher, A12379 |
| Anti-Rabbit Ig 488 | 10 µg/mL | Thermo Fisher, A11034 |
| Anti-Rabbit Ig 594 | 10 µg/mL | Thermo Fisher, Z25307 |
| Anti-Mouse Ig 488 | 10 µg/mL | Thermo Fisher, A21200 |
| Anti-Mouse Ig 568 | 10 µg/mL | Thermo Fisher, A11004 |
| Anti-Guinea Pig Ig 594 | 10 µg/mL | Sigma Aldrich, SAB4600080 |

**Supplementary Table S2.**
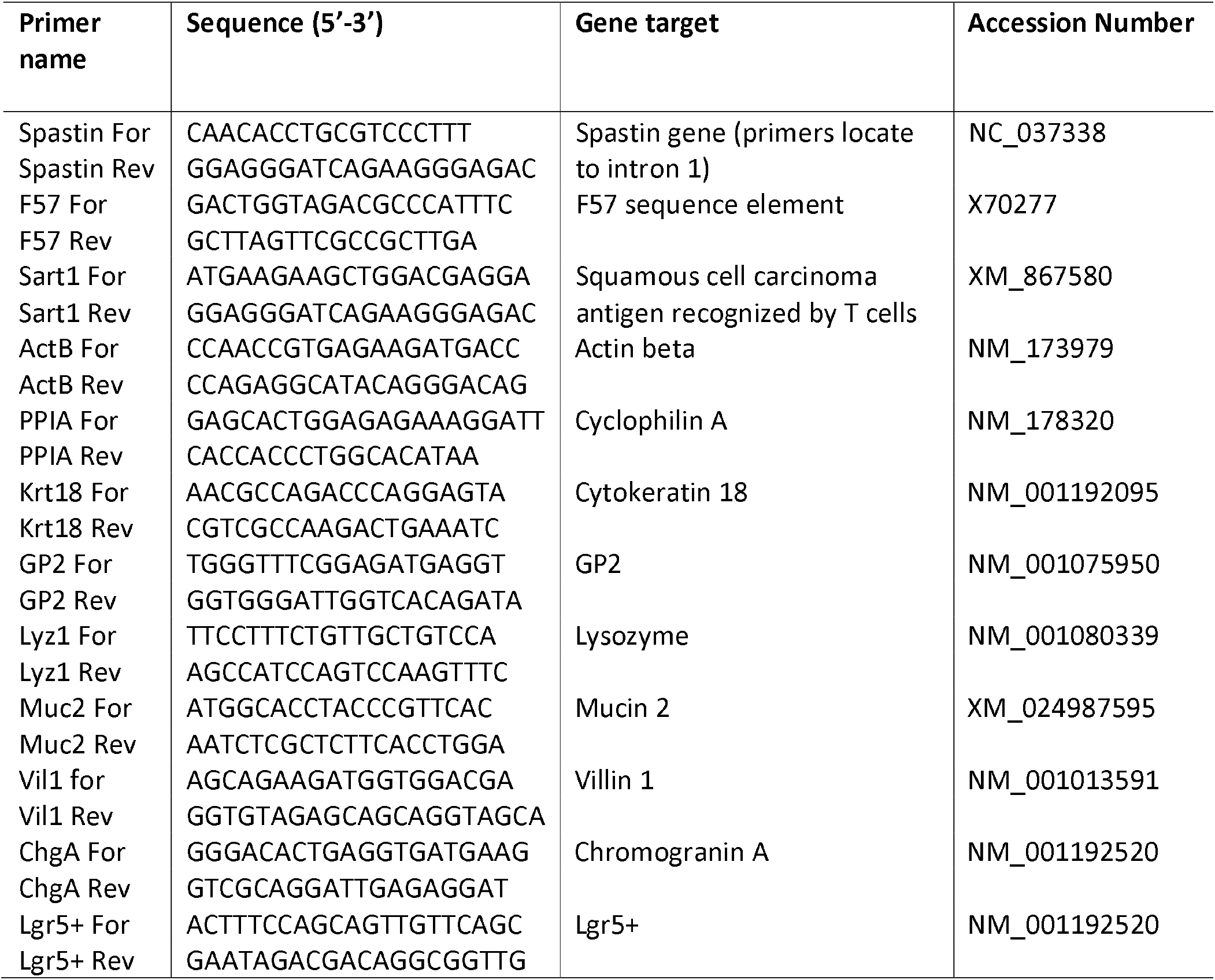
Primer sequences for qRT-PCR experiments

